# MAPK Phosphatase-5 is required for TGF-β signaling through a JNK-dependent pathway

**DOI:** 10.1101/2024.06.27.600976

**Authors:** Sam Dorry, Sravan Perla, Anton M. Bennett

## Abstract

Mitogen-activated protein kinase (MAPK) phosphatases (MKPs) constitute members of the dual-specificity family of protein phosphatases that dephosphorylate the MAPKs. MKP-5 dephosphorylates the stress-responsive MAPKs, p38 MAPK and JNK, and has been shown to promote tissue fibrosis. Here, we provide insight into how MKP-5 regulates the transforming growth factor-β (TGF-β) pathway, a well-established driver of fibrosis. We show that MKP-5-deficient fibroblasts in response to TGF-β are impaired in SMAD2 phosphorylation at canonical and non-canonical sites, nuclear translocation, and transcriptional activation of fibrogenic genes. Consistent with this, pharmacological inhibition of MKP-5 is sufficient to block TGF-β signaling, and that this regulation occurs through a JNK-dependent pathway. By utilizing RNA sequencing and transcriptomic analysis, we identify TGF-β signaling activators regulated by MKP-5 in a JNK-dependent manner, providing mechanistic insight into how MKP-5 promotes TGF-β signaling. This study elucidates a novel mechanism whereby MKP-5-mediated JNK inactivation is required for TGF-β signaling and provides insight into the role of MKP-5 in fibrosis.

## Introduction

Fibrosis is the pathological accumulation of extracellular matrix components in tissues that leads to an impairment of healthy organ function.^1–3^ It is typically driven by dysregulated tissue repair in response to chronic injury and inflammation.^1,4,5^ Fibrosis occurs in many tissues, including the skin, lung, liver, kidney, and skeletal muscle, and is estimated to contribute to approximately 45% of deaths in the industrialized world.^1,6^ Despite the prevalence of fibrosis, treatment options for fibrotic diseases are limited, indicating a need for further development of better antifibrotic therapies.^2,7^

The mitogen activated protein kinases (MAPKs) comprise one of the cell’s most important signaling cascades and drives many essential cellular functions, such as proliferation, differentiation, and regulation of apoptosis.^8–11^ MAPK signaling is necessary to maintain healthy cellular function, so strict regulation of its activity is essential.^8^ Major regulators of the MAPKs are the MAPK phosphatases (MKPs).^12,13^ The MKPs are members of the dual-specificity phosphatase (DUSPs) family of protein phosphatases that are able to dephosphorylate both phospho-tyrosine and phospho-serine/threonine residues.^13^ The MKPs inactivate the MAPKs through specific dephosphorylation of the regulatory phosphotyrosine and phosphothreonine residues within the MAPK activation loop, and as such, represent important regulators of MAPK signaling. MKP-5 is a regulator of the stress-responsive MAPKs, p38 MAPK and c-Jun NH_2_-terminal kinase (JNK).^14,15^ MKP-5 has been shown to contribute to the pathogenesis of fibrosis in multiple tissues.^16–18^ This was first observed when *Mkp5^-/-^* mice were crossed with a mouse model of Duchenne muscular dystrophy (*mdx*).^19,20^ As *mdx* mice age, healthy skeletal muscle is progressively replaced with fibrotic tissue.^21^ However, *mdx* mice lacking MKP-5 had a marked reduction in skeletal muscle fibrosis accompanied by preserved skeletal muscle function demonstrating that MKP-5 is necessary for the progression of fibrosis.^17^ Additionally, we have demonstrated that *Mkp5^-/-^* mice are resistant to bleomycin-induced pulmonary fibrosis, as well as pressure overload-induced cardiac fibrosis.^16,18^. These observations strongly support the interpretation that MKP-5 is required for either the initiation and/or progression of fibrosis in multiple tissues and suggests a fundamental role for MKP-5 in this process.

One of the major mechanisms through which fibrosis occurs is through the transforming growth factor β (TGF-β) pathway.^22,23^ In humans with fibrotic disease, higher levels of TGF-β expression correlate with severity of fibrosis and can predict fibrosis progression.^24^ In animal models, TGF-β overexpression is sufficient to drive fibrosis in multiple tissues.^25–27^ TGF-β signaling regulates the function of many cell types, including macrophages and endothelial cells, but their most prominent contribution to fibrosis is the activation of tissue resident fibroblasts.^22,28,29^ TGF-β promotes proliferation, migration, and cell survival in fibroblasts, and induces expression of collagens and other extracellular matrix proteins.^30,31^ Further, TGF-β can drive the differentiation of fibroblasts to myofibroblasts, a cell type characterized by its expression of fibrogenic genes such as α-smooth muscle actin (*Acta2*), which is a hallmark of scar tissue.^22^ Canonical TGF-β signaling is initiated when the activated TGF-β receptor phosphorylates SMAD2/SMAD3 (R-SMADs).^32–34^ Phosphorylated R-SMADs then complex with SMAD4 and translocate to the nucleus, where this complex binds SMAD-responsive elements in the promoter of genes that are involved in the expression of extracellular matrix proteins.^32–34^ During tissue repair, *Mkp5^-/-^* mice have diminished R-SMAD phosphorylation, as do primary MKP-5-deficient fibroblasts directly stimulated with TGF-β, suggesting that MKP-5 contributes to pro-fibrotic TGF-β signaling.^16,35^ However, the complete mechanisms by which MKP-5 promotes this pathway remain unknown.

In this study, we show that MKP-5 is necessary for TGF-β-mediated SMAD2 phosphorylation at canonical and noncanonical sites, nuclear translocation, and downstream transcription of fibrogenic genes. By utilizing an MKP-5 specific allosteric inhibitor, we show that pharmacological inhibition of MKP-5 recapitulates the effects of MKP-5 genetic deficiency and is also sufficient to block TGF-β signaling. Mechanistically, we demonstrate that MKP-5 regulates TGF-β signaling through a JNK-dependent pathway, and that inhibiting JNK rescues TGF-β-driven SMAD2 phosphorylation and transcription in cells treated with an MKP-5 inhibitor. We utilize RNA sequencing to identify a set of TGF-β activators regulated by MKP-5/JNK, further elucidating the molecular mechanisms of regulation. This study elucidates a novel mechanism whereby MKP-5-mediated JNK inactivation is required for TGF-β signaling and provides insight into the role of MKP-5 in fibrosis.

## Results

### MKP-5 promotes TGF-β-signaling and SMAD2 phosphorylation

To characterize the involvement of MKP-5 in TGF-β signaling, we first sought to determine if MKP-5 expression was altered in response to TGF-β stimulation. To investigate this, we stimulated mouse embryonic fibroblasts with TGF-β1 at several time points, ranging from 2 to 24 hr, and measured MKP-5 (*Dusp10*) mRNA expression using qPCR. We found that TGF-β stimulation induces an acute upregulation of *Dusp10* mRNA at 2 hr, followed by downregulation at later time points (Figure 1A). Considering our previous observations that MKP-5 is necessary for TGF-β signaling, this data suggests a previously uncharacterized axis of mutual regulation between TGF-β and MKP-5/MAPK signaling.^16^

**Figure 1.**
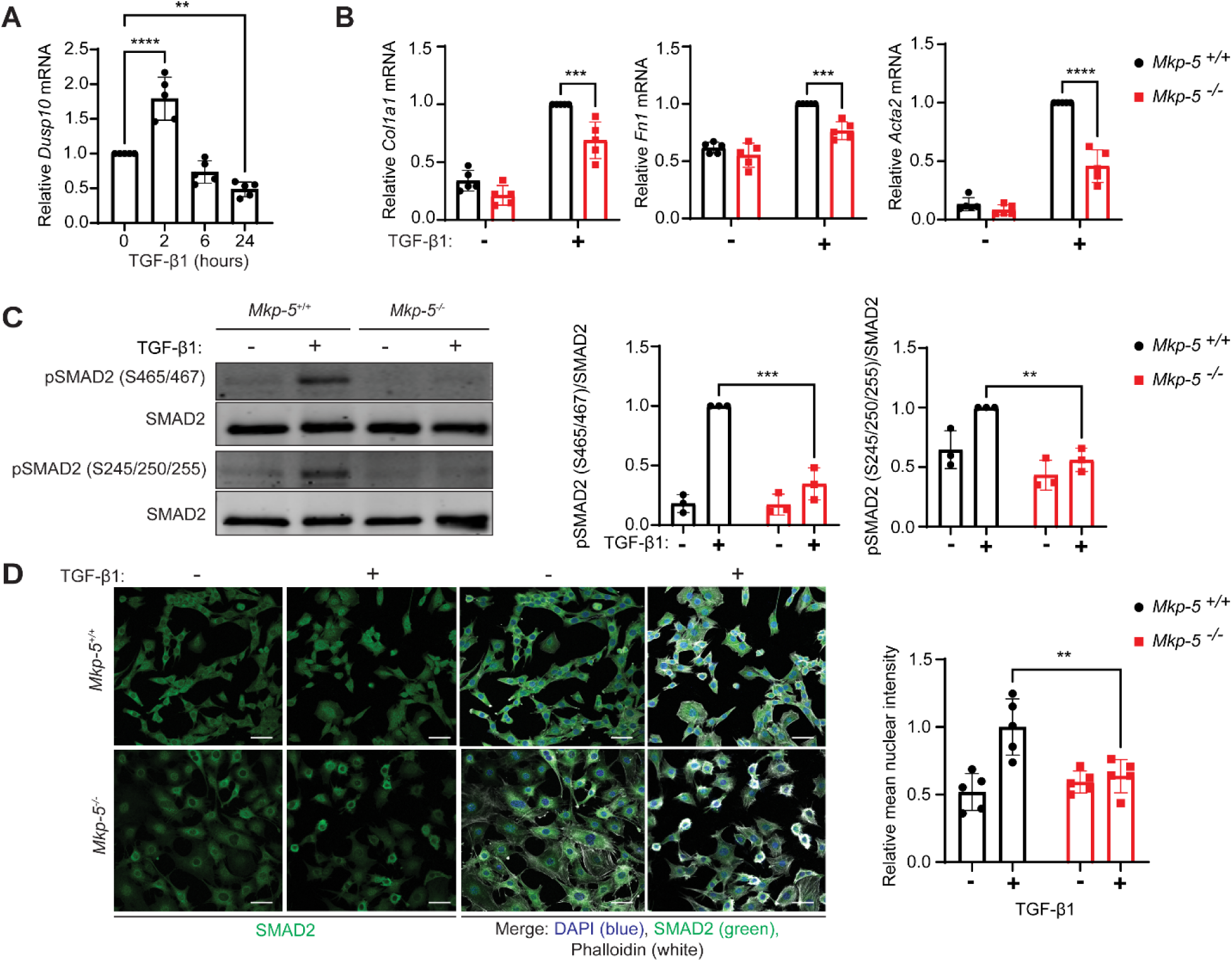
MKP-5 promotes TGF-β signaling in fibroblasts. (A) Wild-type mouse embryonic fibroblasts were stimulated with TGF-β1 (2 ng/mL) at the indicated time points, and expression of MKP-5 (*Dusp10)* was measured by qPCR. (B) *Mkp-5^+/+^* and *Mkp-5^-/-^* fibroblasts were stimulated with TGF-β1 (2 ng/mL) for 24 hr, and expression levels of *Col1a1, Fn1,* and *Acta2* were measured by qPCR. Data from (A) and (B) represent the mean ± SEM derived from 5 independent experiments. (C) *Mkp-5^+/+^* and *Mkp-5^-/-^* fibroblasts were stimulated with TGF-β1 (2 ng/mL) for 10 minutes, and SMAD2 phosphorylation was measured using anti-pSMAD2 (S465/467), anti-pSMAD2 (S245/250/255) and anti-SMAD2 antibodies. Right panel represents the densitometry of immunoblots from pSMAD2 (S465/467) and pSMAD2 (S245/S250/255) normalized to total SMAD2. Data represent the mean ± SEM derived from 3 independent experiments. (D) *Mkp-5^+/+^* and *Mkp-5^-/-^* fibroblasts were stimulated with TGF-β1 (2 ng/mL) for 10 minutes, fixed, and stained with anti-SMAD2 (green), phalloidin (white), and DAPI (blue). Scale bar = 50 μM. Images were analyzed and nuclear intensity of SMAD2 was quantified using ImageJ. Key: *\*\*: p*-value<0.01, ***: *p*-value<0.001, ****: *p*-value<0.0001.

We sought to characterize the role of MKP-5 in regulating specific components within the TGF-β signaling pathway. To investigate the impact of MKP-5 on TGF-β-target gene expression, we stimulated *Mkp5^+/+^* and *Mkp5^-/-^* fibroblasts with TGF-β1 for 24 hr and measured expression of TGF-β target genes using qPCR. We found that *Mkp5^-/-^* fibroblasts in response to TGF-β1 had significantly diminished expression of collagen (*Col1a1*), fibrobectin (*Fn1*), and actin (*Acta2*) as compared with *Mkp5^+/+^* treated cells. These results indicate that MKP-5 is necessary for TGF-β-mediated transcription of established fibrotic genes (Figure 1B). We also investigated the role of MKP-5 in regulating TGF-β-driven SMAD2 phosphorylation. In canonical TGF-β signaling, the activated TGF-β receptor directly phosphorylates the c-terminus site of SMAD2 (S465/467), which induces a conformational change that allows SMAD2 to complex with SMAD4, translocate to the nucleus, and facilitate transcription of TGF-β target genes.^32–34^ Additionally, the TGF-β receptor also activates various non-canonical signaling pathways, including the MAPK, Rho kinase, and phosphatidylinositol 3’-kinase/AKT kinases.^34,36^ These pathways can converge on the TGF-β/SMAD signaling by phosphorylating SMAD2 on its linker site (S245/250/255).^36,37^ The effect of SMAD2 linker phosphorylation on its activity is complex and has been reported to both promote and inhibit transcription depending on the cellular context.^37–39^ To determine the effect of MKP-5 on SMAD2 phosphorylation, *Mkp5^+/+^* and *Mkp5^-/-^* fibroblasts were stimulated with TGF-β1 for 10 minutes, and protein was isolated for immunoblotting. As expected, TGF-β1-stimulated *Mkp5^+/+^* fibroblasts exhibited a robust induction of SMAD2 c-terminus and linker phosphorylation (Figure 1C). In contrast, *Mkp5^-/-^* fibroblasts were shown to have significantly diminished TGF-β1-induced phosphorylation of SMAD2 on both the c-terminus and linker sites (Figure 1C). SMAD2 c-terminus phosphorylation is required for its complex formation with SMAD4 and nuclear translocation. Therefore, we tested whether abrogating MKP-5 expression also resulted in impaired TGF-β-driven SMAD2 nuclear localization. *Mkp5^+/+^* and *Mkp5^-/-^* fibroblasts were stimulated with TGF-β1 for 10 minutes, then cells were fixed and stained with anti-SMAD2 antibody, phalloidin, and DAPI (Figure 1D). Images were collected using confocal microscopy and analyzed using ImageJ software. The average fluorescence intensity derived from SMAD2 staining was quantified within the nucleus of cells and compared between groups (Figure 1D). In response to TGF-β1 stimulation, *Mkp5^-/-^* fibroblasts had significantly diminished SMAD2 nuclear accumulation compared with *Mkp5^+/+^* fibroblasts (Figure 1D). Collectively, these results demonstrate that MKP-5 is required for both canonical and non-canonical TGF-β signaling and subsequent downstream translocation of the activated SMAD complex to the nucleus.

### MKP-5 activity is required for TGF-β signaling

Previously, a high throughput screen identified a novel allosteric MKP-5 inhibitor (Cmpd 1). Cmpd 1 was found to upregulate both p38 MAPK and JNK activities in cells, promote myoblast differentiation and diminish TGF-β-induced SMAD2 phosphorylation.^35^ Additional MKP-5 allosteric inhibitors have been developed that either show equipotency or improved potency as compared with Cmpd 1.^40^ To more effectively interrogate the role of MKP-5 in TGF-β signaling, we utilized a derivative of Cmpd 1, referred to herein as Cmpd 2, which is more potent than Cmpd 1 and maintains MKP-5 selectivity (Supplementary material, Table S1). MKP-5 preferentially dephosphorylates the stress-responsive MAPKs, p38 MAPK and JNK.^12^ Consistent with this MKP-5-deficient fibroblasts demonstrate hyperactivation of p38 MAPK and JNK, but not ERK.^35^ To test the efficacy of Cmpd 2, we incubated 3T3 fibroblasts with Cmpd 2 (10 µM) for 4 hr, and measured levels of phosphorylated MAPKs. Under these conditions, we observed that Cmpd 2 significantly increased phosphorylation of JNK and p38 MAPK, by 1.5- and 1.4-fold, respectively relative to DMSO-treated fibroblasts (Figure 2A). Importantly, Cmpd 2 did not induce an upregulation of ERK phosphorylation (Figure 2A) and Cmpd 2 was shown to not effect either fibroblast proliferation or cytotoxicity at concentrations up to 30 µM (Supplementary Material, Figure S1). These results are consistent with the expected effects of selective MKP-5 catalytic inhibition.^35^

**Figure 2.**
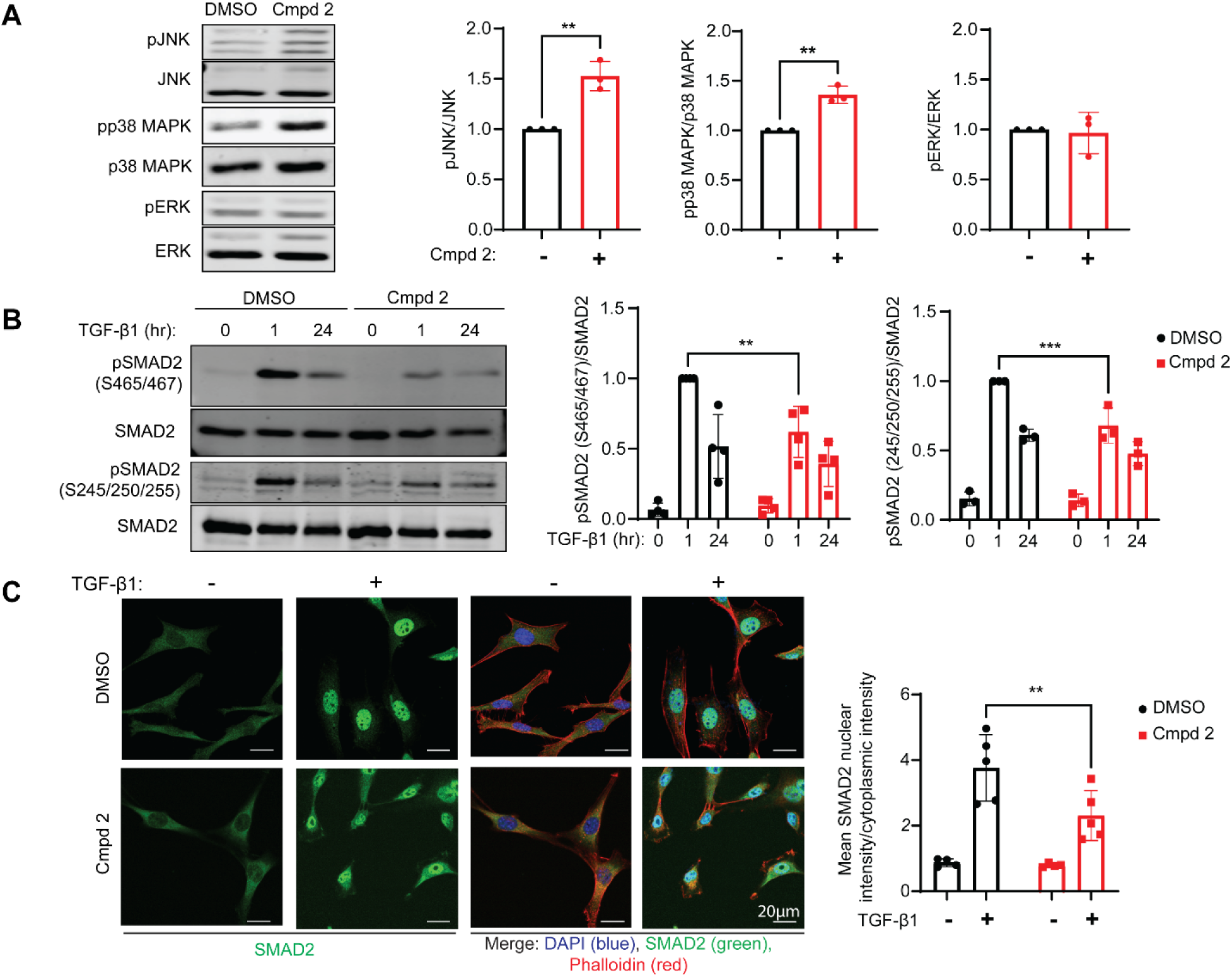
MKP-5 activity is required for TGF-β-mediated SMAD2 phosphorylation and nuclear translocation. (A) 3T3 fibroblasts were treated with Cmpd 2 (10 µM) for 4 hr. Shown are immunoblots for pJNK, pp38 MAPK and pERK1/2 and respective MAPK controls. Right panel shows densitometry from 3 independent experiments representing the ratios of pMAPK/MAPK. (B) 3T3 fibroblasts treated with Cmpd 2 (10 µM) for 4 hr and stimulated with TGF-β1 (2 ng/mL) for either 1 or 24 hr were immunoblotted for SMAD2 phosphorylation using anti-pSMAD2 (S465/467), anti-pSMAD2 (S245/250/255) and anti-SMAD2 antibodies. Right panel shows the densitometry of immunoblots from pSMAD2 (S465/467) and pSMAD2 (S245/S250/255) normalized to total SMAD2. Data represent the mean ± SEM derived from 4 independent experiments. (C) 3T3 fibroblasts treated with Cmpd 2 (10 µM) for 4 hr were stimulated with TGF-β1 (2 ng/mL) for 1 hr. Cells were fixed and stained with anti-SMAD2 (green), phalloidin (red), and DAPI (blue). Images were analyzed and SMAD2 localization was quantitated using ImageJ. Key: *\*\*: p*-value<0.01, ***: *p*-value<0.001, ****: *p*-value<0.0001.

We next tested the effect of MKP-5 inhibition with Cmpd 2 on TGF-β-mediated SMAD2 phosphorylation. 3T3 fibroblasts were incubated with DMSO 2 or Cmpd for 4 hr, and then stimulated with TGF-β1 at the indicated time points. SMAD2 phosphorylation was determined by immunoblotting. Cmpd 2-treated 3T3 fibroblasts had significantly reduced levels of SMAD2 phosphorylation on both the c-terminus and linker region sites as compared with DMSO-treated cells following 1 hr TGF-β stimulation. However, by 24 hr of TGF-β treatment, Cmpd 2-treated cells showed equivalent levels of phosphorylation of SMAD2 on both the c-terminus and linker region as compared with DMSO-treated cells (Figure 2B). These results show that Cmpd 2 can acutely inhibit TGF-β-mediated SMAD2 phosphorylation on both canonical and noncanonical sites. We next investigated the impact of MKP-5 inhibition by Cmpd 2 on TGF-β-mediated SMAD2 nuclear translocation, which occurs as a result of SMAD2 c-terminus phosphorylation. We incubated 3T3 fibroblasts with either DMSO or Cmpd 2 (10 µM) for 4 hr, followed by stimulation with TGF-β1 for 1 hr and SMAD2 sub-cellular localization was assessed. We found that Cmpd 2-treated fibroblasts accumulated significantly less nuclear SMAD2 compared with DMSO-treated cells, indicating that MKP-5 catalytic inhibition prevents TGF-β-mediated SMAD2 nuclear translocation (Figure 2D).

To assess the role of MKP-5 catalysis on TGF-β-mediated transcription, we treated 3T3 fibroblasts with DMSO or Cmpd 2 (10 µM) and stimulated cells with TGF-β1 for 24 hr. Expression of known TGF-β-target genes *Col1a1*, *Fn1*, and *Acta2* were measured using qPCR. While fibroblasts treated with TGF-β showed increased expression levels of *Col1a1*, *Fn1*, and *Acta2* genes, Cmpd 2-treated fibroblasts were significantly inhibited in the expression of these genes indicating that MKP-5 activity is necessary for TGF-β-mediated transcription (Figure 3A). Notably, Cmpd 2 did not affect expression of these genes in *Mkp5^-/-^* fibroblasts, validating that Cmpd 2 is regulating TGF-β signaling through an on-target, MKP-5-mediated mechanism (Supplemental material, Figure S2). In addition to measuring the expression of established TGF-β target genes, we complemented these experiments using a TGF-β-responsive luciferase assay that specifically reports activity through SMAD complex binding to its cognate SMAD-binding-element (SBE). A549 cells were stably transfected with a luciferase reporter containing four SBE repeats (SBE reporter cells). In these cells, TGF-β stimulation induces SMAD-dependent transcription of luciferase (Figure 3B). SBE reporter cells were treated with either DMSO or Cmpd 2 and stimulated with TGF-β overnight. Here we found that DMSO-treated SBE reporter cells stimulated with TGF-β induced a robust induction of SBE-mediated luciferase activity (Figure 3B). In contrast, TGF-β-stimulated SBE reporter cells treated with Cmpd 2 exhibited a significantly reduced level of SBE-mediated luciferase activity as compared with DMSO-treated cells (Figure 3B). These data indicate that inhibition of MKP-5 activity specifically blocks SMAD-driven transcriptional activation in response to TGF-β. In addition, Cmpd 2 exhibited a dose-dependent inhibition of SBE-mediated luciferase activity with an EC_50_ of 0.9 µM (Figure 3C). In addition to regulating transcription, we also determined if Cmpd 2 inhibits expression of α-SMA protein in response to TGF-β. By incubating 3T3 fibroblasts with either DMSO or Cmpd 2 for 4 hr, followed by stimulation with TGF-β for either 24 or 48 hr, we found that Cmpd 2-treated cells had diminished expression of α-SMA protein at 24 hr (Figure 3D). Collectively, these results demonstrate that MKP-5 activity is required to promote TGF-β-mediated SMAD2 phosphorylation, nuclear translocation and transcriptional activation.

**Figure 3.**
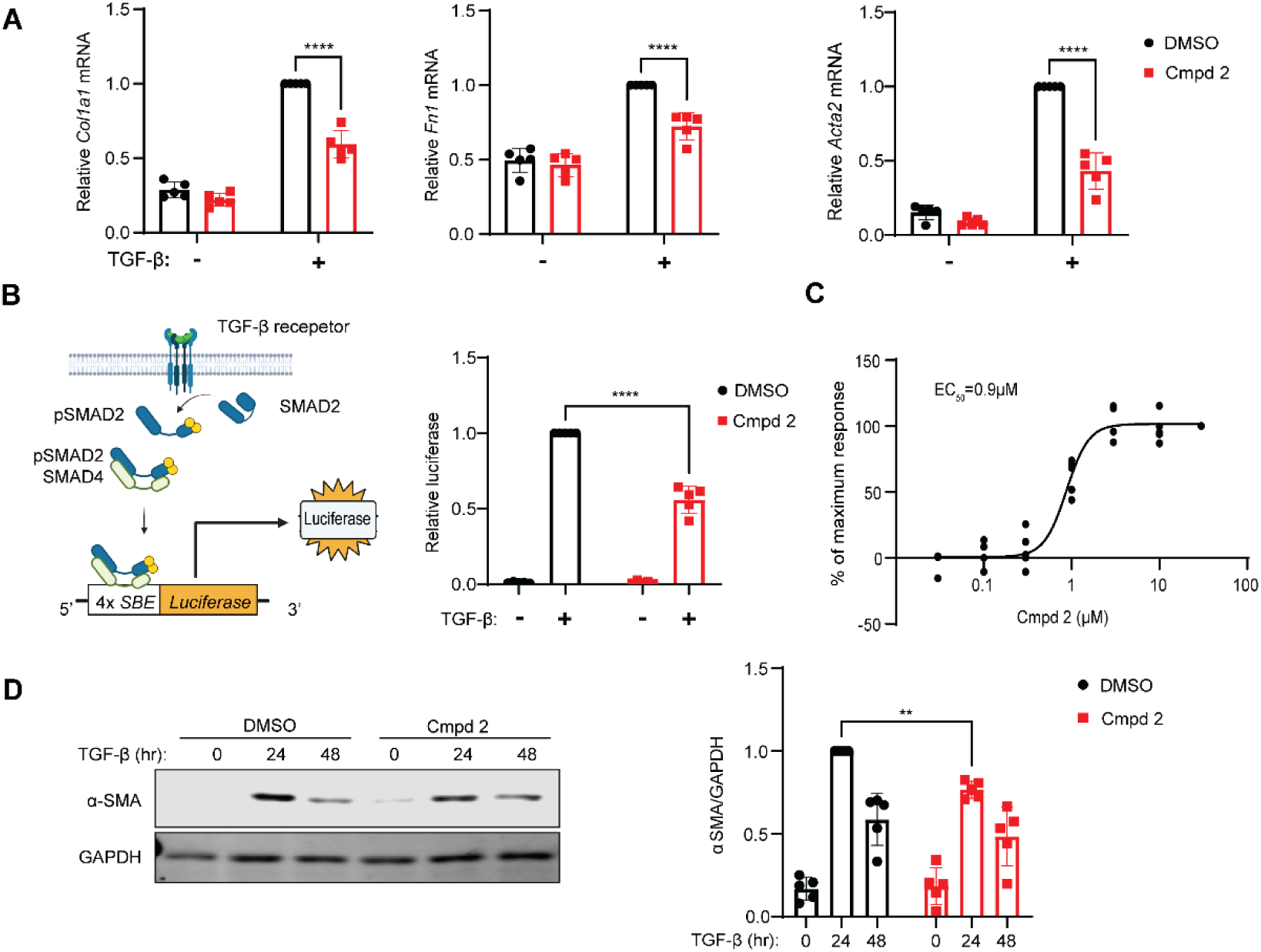
MKP-5 activity is required for TGF-β-mediated gene expression. (A) 3T3 fibroblasts were treated with Cmpd 2 (10 µM) and TGF-β1 (2 ng/mL) for 24 hr, expression of TGF-β target genes *Col1a1, Fn1,* and *Acta2* were measured using qPCR. Data represent the mean ± SEM derived from 5 independent experiments. (B) Left panel represents a schematic of the SMAD binding element (SBE) stable cell line reporter system. A549 cells were stably transfected with four SBE repeats (4xSBE) fused to a luciferase reporter (SBE reporter cells). Right panel shows SBE reporter cells treated with Cmpd 2 (10 µM) and TGF-β1 (2 ng/mL) for 24 hr and luciferase activity measured. (C) SBE reporter cells were treated with TGF-β1 (2 ng/mL) and various concentrations of Cmpd 2 for 24 hr and luciferase activity measured. (D) 3T3 fibroblasts were treated with Cmpd 2 (10 µM) for 4 hr, then stimulated with TGF-β1 (2 ng/mL) for either 24 or 48 hr and immunoblotted using anti-α-SMA and GAPDH antibodies. Densitometry values are shown on the right representing the mean ± SEM from 5 independent experiments. Key: *\*\*: p*-value<0.01, ***: *p*-value<0.001, ****: *p*-value<0.0001.

### MKP-5 regulates TGF-β signaling through a JNK-dependent mechanism

MKP-5 negatively regulates the activity of p38 MAPK and JNK.^12^ In cells lacking MKP-5, the enhanced activity of either p38 MAPK and/or JNK is likely to participate in the inhibition of TGF-β signaling. Thus, inhibition of either p38 MAPK and/or JNK in context of MKP-5 deficiency would be anticipated to rescue TGF-β signaling. To test this, we measured TGF-β signaling in SBE reporter cells treated with Cmpd 2 and various MAPK inhibitors. Cells were stimulated with TGF-β and incubated with Cmpd 2 in the presence of either DMSO or 10 µM of U0126 (ERK inhibitor), SB203580 (p38 MAPK inhibitor) or 10 µM SP600125 (JNK inhibitor) overnight (Figure 4A). SBE reporter cells treated with Cmpd 2 induced ∼50% SBE-mediated luciferase activity as compared with DMSO-treated cells. U0126-treated cells did not significantly affect SBE-mediated luciferase activity in either DMSO or Cmpd 2-treated cells. SB203580 reduced SBE-mediated luciferase activity in both DMSO and Cmpd 2-treated cells. Remarkably, cells treated with both Cmpd 2 and SP600125 had similar levels of SBE-mediated luciferase activity compared with DMSO-treated cells. These results demonstrate that inhibiting JNK, but neither ERK nor p38 MAPK, results in the rescue of TGF-β signaling in Cmpd 2-treated cells. Thus, suggesting that the effect of MKP-5 in promoting TGF-β signaling is dependent upon JNK dephosphorylation.

**Figure 4.**
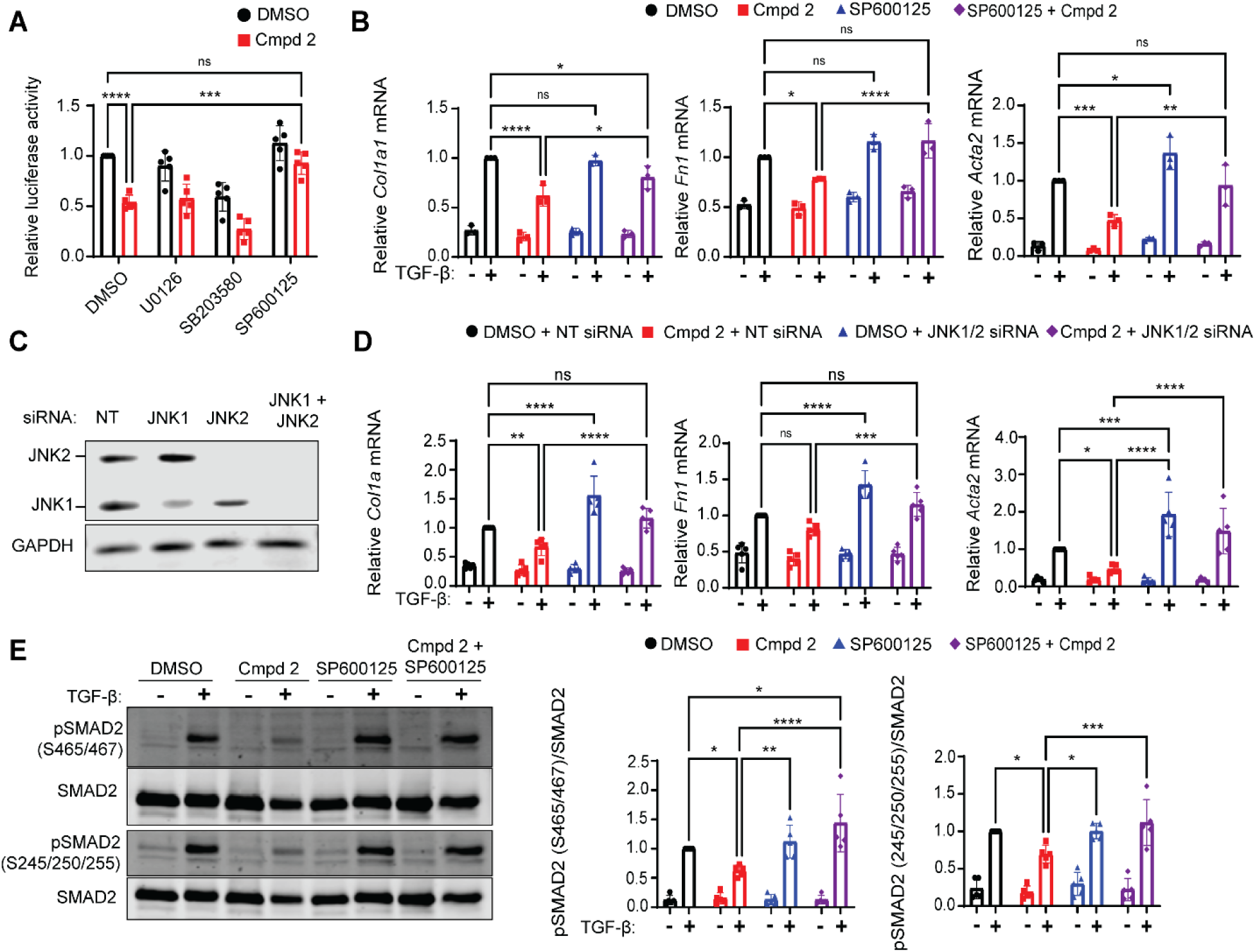
MKP-5 regulates TGF-β signaling through a JNK-dependent mechanism. (A) SBE reporter cells were treated with TGF-β1 (2 ng/mL), Cmpd 2 (10 µM), and MAPK inhibitors U0126, SB203580 and SP600125 at 10 µM for 24 hr. Cells were lysed and luciferase activity measured. (B) 3T3 fibroblasts were treated with TGF-β1 (2 ng/mL), Cmpd 2 (10 µM), and SP600125 (10 µM) for 24 hr. Expression levels of TGF-β target genes *Col1a1, Fn1,* and *Acta2* were measured by qPCR. (C) 3T3 fibroblasts were treated with siRNA’s to JNK1, JNK2 and JNK1/JNK2. Immunoblots were performed using anti-JNK antibodies. (D) 3T3 fibroblasts were transfected with either non-targeting siRNA (NT siRNA) or JNK1 and JNK2 siRNA (JNK1/2 siRNA) for 48 hr, and cells were treated with Cmpd 2 (10 µM) and TGF-β1 (2 ng/mL) for 24 hr expression of TGF-β target genes *Col1a1, Fn1,* and *Acta2* were measured by qPCR. (E) 3T3 fibroblasts were treated with Cmpd 2 (10 µM) and SP600125 (10 µM) for 4 hr, then stimulated with TGF-β1 (2 ng/mL) for 1 hr. Cells were lysed and immunoblotted for SMAD2 phosphorylation using anti-pSMAD2 (S465/467), anti-pSMAD2 (S245/s50/255) and anti-SMAD2 antibodies. Right panel shows the densitometry of immunoblots from pSMAD2 (S465/467) and pSMAD2 (S245/S250/255) normalized to total SMAD2. Data represent the mean ± SEM derived from 5 independent experiments. Key: *\*\*: p*-value<0.01, ***: *p*-value<0.001, ****: *p*-value<0.0001.

To investigate the involvement of MKP-5 in regulating expression of TGF-β target genes through JNK, 3T3 fibroblasts were stimulated with TGF-β1 and treated with Cmpd 2 and/or SP600125 overnight. Expression of *Col1a1*, *Fn1*, and *Acta2* were measured using qPCR (Figure 4B). Cells treated with Cmpd 2 had significantly diminished expression of *Col1a1*, *Fn1*, and *Acta2* relative to DMSO-treated fibroblasts. In contrast, fibroblasts treated with both Cmpd 2 and SP600125 had significantly increased expression of *Col1a1*, *Fn1*, and *Acta2* compared with fibroblasts treated with Cmpd 2 alone, indicating that JNK inhibition in context of MKP-5 impaired activity rescues TGF-β signaling (Figure 4B). Notably, inhibition of p38 MAPK activity failed to rescue expression of TGF-β-target genes in Cmpd 2-treated cells (Supplemental material, Figure S3). To substantiate the involvement of MKP-5 in regulating expression of TGF-β target genes through JNK, we tested the effect of knocking down JNK1/2 on TGF-β signaling in Cmpd 2-treated cells. First, we transfected 3T3 fibroblasts with either non-targeting (NT), JNK1, JNK2 or JNK1 and JNK2 (JNK1/2) siRNAs for 48 hr. The expression of JNK1 and JNK2 was determined by immunoblotting and these results showed effective knockdown of both JNK1 and JNK2 alone as well as JNK1 and JNK2 combined in 3T3 fibroblasts (Figure 4C). Next, fibroblasts were transfected with either NT or JNK1 plus JNK2 siRNA (JNK1/2) for 48 hr, followed by treatment with Cmpd 2 and TGF-β1 overnight (Figure 4D). When TGF-β1 target genes, *Col1a1*, *Fn1*, and *Acta2*, were measured by qPCR we found that JNK1/2 siRNA-treated fibroblasts exhibited enhanced gene expression as compared with

DMSO-treated cells (Figure 4D). Cmpd 2-treated cells, as expected had decreased *Col1a1*, *Fn1*, and *Acta2* gene expression levels in response to TGF-β1 but the expression levels of these genes were restored to that of controls in fibroblasts treated with Cmpd 2 plus JNK1/2 siRNA (Figure 4D). These results confirm those shown in Figure 4B and substantiate the interpretation that MKP-5 mediates its effects on TGF-β signaling through JNK. Finally, we tested the role of JNK in MKP-5-medaited regulation of SMAD2 phosphorylation. 3T3 fibroblasts were treated with Cmpd 2 and/or SP600125 for 4 hr, then fibroblasts were stimulated with TGF-β1 for 1 hr. In response to TGF-β stimulation, Cmpd 2-treated cells had diminished SMAD2 phosphorylation at both the c-terminus and linker sites compared with DMSO-treated fibroblasts (Figure 4E). TGF-β-stimulated fibroblasts treated with both Cmpd 2 and SP600125 exhibited levels of SMAD2 phosphorylation that were equivalent to fibroblasts treated with DMSO control (Figure 4E). These results indicate that JNK inhibition rescues TGF-β-indued SMAD phosphorylation and gene expression in fibroblasts exhibiting a deficiency in MKP-5 activity. The ability to rescue TGF-β-induced signaling in context of MKP-5 inhibition supports the notion that JNK is a major mediator of MKP-5-dependent TGF-β signaling.

### Transcriptome analysis identifies MKP-5-mediated TGF-β regulatory genes

We next sought to elucidate the molecular mechanism by which MKP-5-mediated JNK activity regulates TGF-β signaling. We hypothesized that MKP-5-mediated JNK activity regulates the transcription of either TGF-β signaling activators and/or inhibitors. To identify potential candidate genes, we performed RNA sequencing and transcriptomic analysis of fibroblasts treated with Cmpd 2 and siRNA knockdown of JNK1 and JNK2. We reasoned that relevant Cmpd 2-inhibited/activated genes should be rescued following JNK1/2 knockdown as illustrated in Figure 4. We performed RNA sequencing on 3T3 fibroblasts in three experimental groups. In the first group, cells were incubated in DMSO alone, the second group cells were treated with 10 µM Cmpd 2 (CMPD2), and in the third group cells were treated with 10 µM Cmpd 2 plus siRNA knockdown of JNK1/2 (CMPD2+JNKsiRNA). All experimental groups were stimulated with TGF-β (2 ng/mL) for 24 hr. Candidate genes were identified as genes significantly upregulated or downregulated in CMPD2 relative to DMSO but rescued to levels in CMPD2+JNKsiRNA. Principal component analysis (PCA) showed that samples from each treatment group clustered together (Supplemental material, Figure S4). Gene counts were normalized, and differentially expressed genes (DEGs) (*p* ≤ 0.05, fold-change -1.2≤ 1 ≤ 1.2) were calculated using DESeq2. To identify transcriptional changes driven by catalytic inhibition of MKP-5, we compared DEGs between CMPD2 and DMSO treated groups. CMPD2 generated 64 upregulated and 75 downregulated genes relative to DMSO (Figure 5A, left). To identify MKP-5-mediated JNK1/2 transcriptional targets we compared DEGs between CMPD2+JNKsiRNA and CMPD2 treatment groups which yielded 191 upregulated and 242 downregulated genes, relative to CMPD2 (Figure 5A, right). To identify physiological/pathogenic signaling pathways associated with the TGF-β-induced MKP-5-JNK signaling axis, DEGs were analyzed with Ingenuity Pathway Analysis (IPA) software. CMPD2-treated cells demonstrated inhibition of several signaling pathways associated with fibrosis, namely hepatic fibrosis, idiopathic pulmonary fibrosis, and wound healing (Figure 5B). Specific genes associated with inhibition of hepatic fibrosis and pulmonary fibrosis were identified (Figure 5C and Supplemental material, Table S2). IPA Upstream Regulator analysis further validated that that Cmpd 2-treatment blocked TGF-β signaling in fibroblasts (CMPD2 vs DMSO; z-score = -4.126, *p* =9.60*10^-11^), and that JNK knockdown rescued TGF-β signaling in Cmpd 2-treated cells (CMPD2+JNKsiRNA vs CMPD2; z-score=2.885, *p* =1.97*10^-25^).

**Figure 5.**
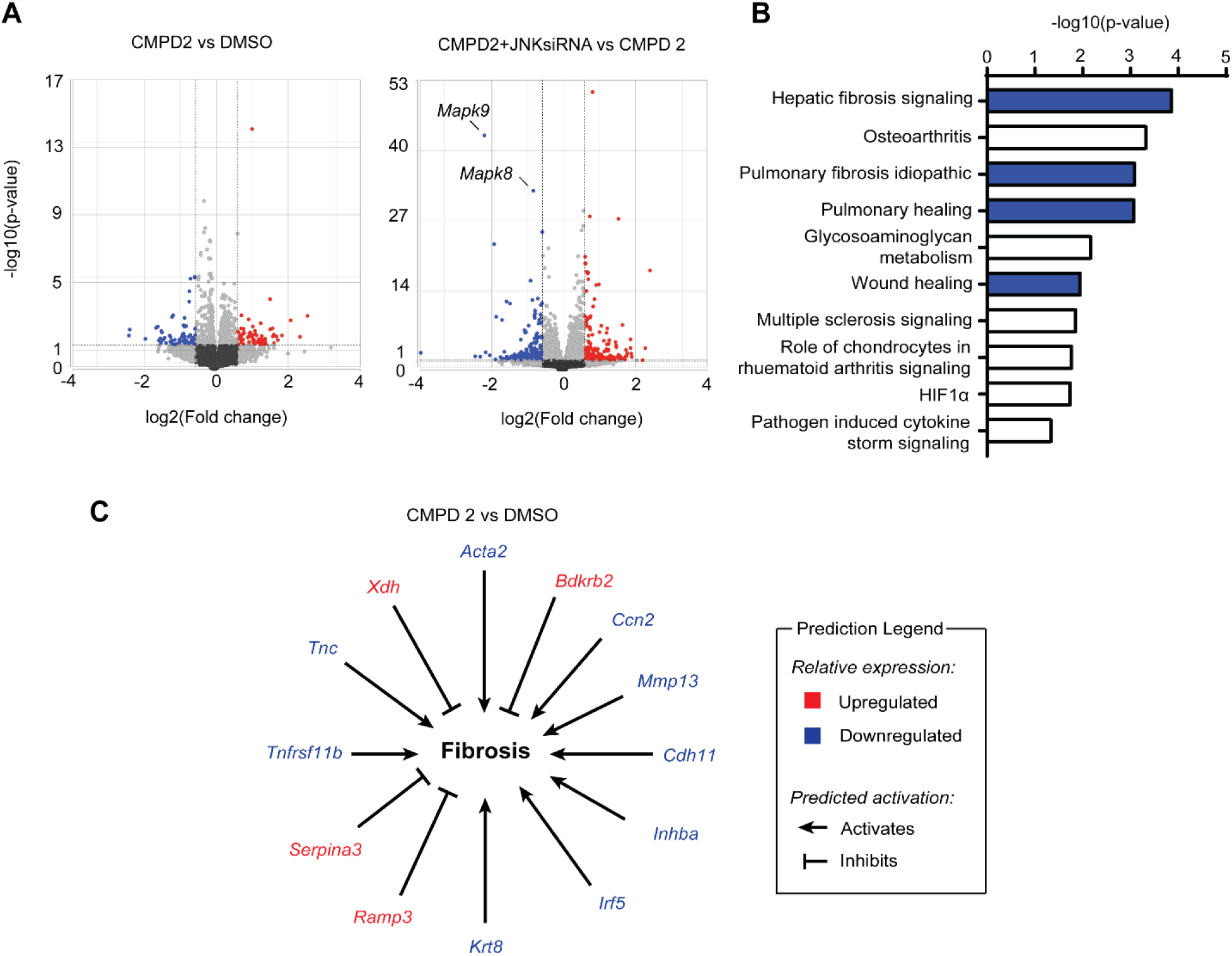
TGF-β-mediated transcription and tissue fibrosis pathways are regulated in a MKP-5-JNK-dependent manner. 3T3 fibroblasts were treated with either non-targeting siRNA (NT siRNA) and DMSO (DMSO), NT siRNA and Cmpd 2 (CMPD2), or JNK1/2-targeting siRNA (JNK siRNA) and Cmpd 2 (CMPD2+JNKsiRNA). All experimental groups were treated with TGF-β1 (2 ng/mL) for 24 hr. (A) Differentially expressed genes (DEGs) of CMPD2 vs DMSO groups and CMPD2+JNKsiRNA vs CMPD2 groups displayed as volcano plots. Cut-off values of *p ≤* 0.05 and fold change ≥ 1.5 (red) or ≤ -1.5 (blue) were used to identify DEGs. (B) Ingenuity Pathway Analysis showing the top ten downregulated (z-score) signaling pathways by Cmpd 2-treatment. Pathways associated with fibrosis are highlighted in blue. (C) Schematic representation showing the effect of Cmpd 2 on expression of genes associated with fibrosis.

We also performed Gene Set Enrichment Analysis (GSEA) using the Mouse Molecular Signatures Database.^41,42^ The analysis compared expression values for CMPD2 to the pooled expression values for DMSO and CMPD2+JNKsiRNA (REST). Using the M2 curated gene set analysis, we identified that the gene set most significantly altered between CMPD2 and REST corresponded to expression of TGF-β target genes (KARLSSON_TGFb1_TARGETS_UP, ES=-0.6300272, *p* <0.001) (Figure 6A and B). This indicates Cmpd 2 inhibits expression of TGF-β-target genes relative to DMSO-treated cells, and that JNK knockdown rescues expression of those genes in Cmpd 2-treated cells. We also attempted to identify MKP-5-JNK-regulated genes that could mediate the observed changes in TGF-β signaling. To determine candidate genes, we identified targets that were either upregulated or downregulated by Cmpd 2-treatment (CMPD2 vs DMSO; 0.80 ≥ fold change ≥ 1.2, *p* ≤ 0.05) and rescued by JNK1/2 knockdown (CMPD2+JNKsiRNA vs CMPD2; 0.80 ≥ fold change ≥ 1.2, *p* ≤ 0.05) (Figure 6C). We identified a total of 305 DEGs, 137 downregulated and 168 upregulated candidate genes (Figure 6C). To substantiate the validity of this non-biased screen, we validated the expression of identified target genes known to promote TGF-β signaling. As described for the RNA sequencing experiments, we assigned three treatment groups representing DMSO control, Cmpd 2-treated and Cmpd 2 plus JNK1/2 siRNA-treated fibroblasts. Cells were treated with TGF-β for 24 hr followed by qPCR analysis for the expression of target genes. One of the most well-established targets genes in the TGF-β pathway is *Acta2* which we showed was downregulated in Cmpd 2-treated fibroblasts and rescued in Cmpd 2 plus JNK1/2 siRNA-treated fibroblasts (Figure 6D). This result confirmed the successful setup of our experimental scheme to identify relevant TGF-β-responsive genes regulated via the MKP-5-JNK signaling axis. We next validated the following highly significant gene candidates; *Thbs1, Inhba, Nuak2,* and *Cdh11* which have been shown to be involved in the activation of TGF-β signaling.^43–45^ Expression levels of these TGF-β signaling activators were inhibited by Cmpd 2-treatment and rescued by JNK knockdown (Figure 6D). These results suggest that *Thbs1, Inhba, Nuak2,* and *Cdh11* genes may mediate MKP-5-JNK regulation of TGF-β signaling. Collectively, our transcriptomic analysis is in line with the interpretation that MKP-5 activity in TGF-β signaling is involved in the regulation of fibrosis consistent with our previous findings that show MKP-5-deficient mice are resistant to the development of muscle, lung and cardiac fibrosis.^16,18,46^

**Figure 6.**
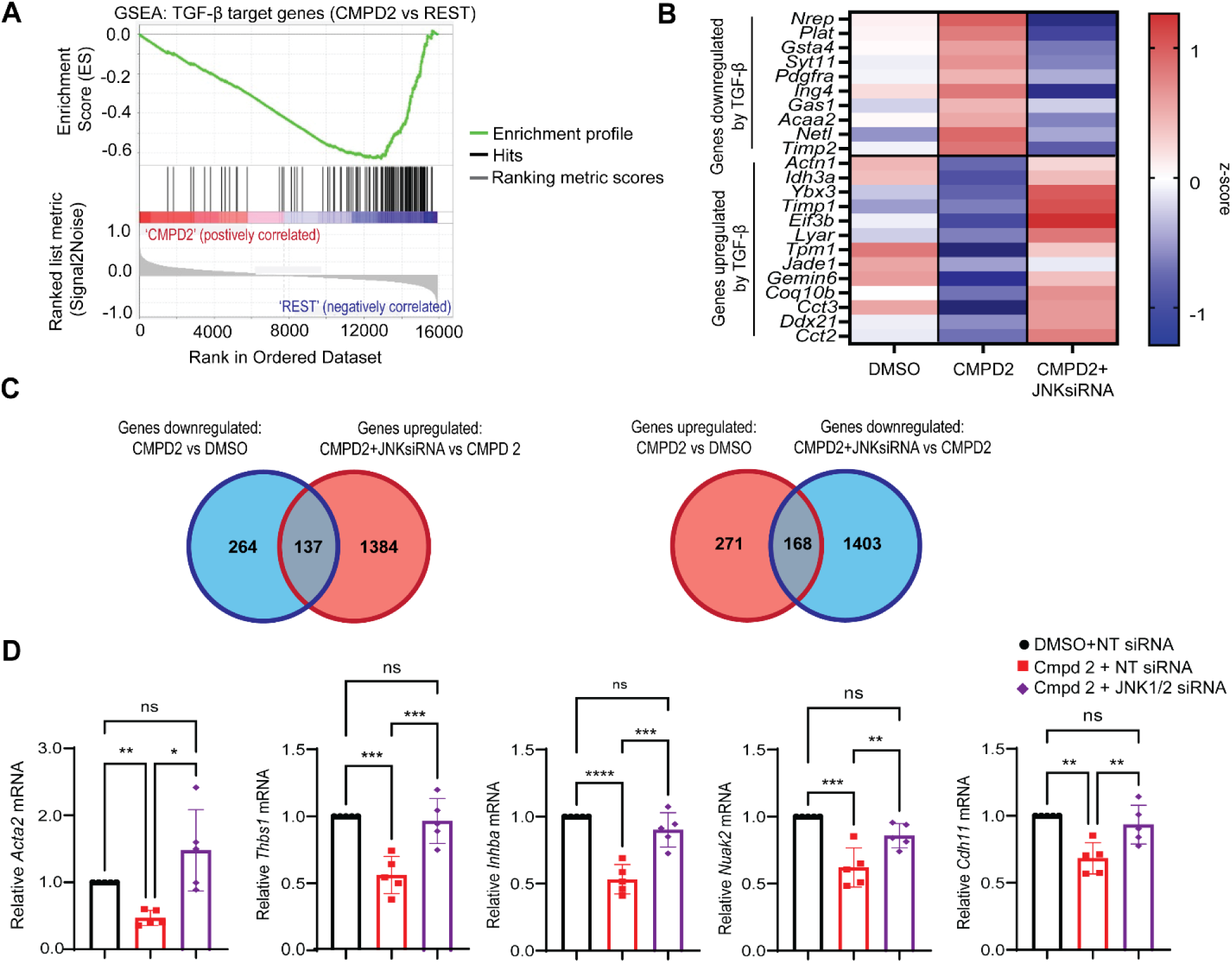
Identification of MKP-5-JNK-dependent TGF-β regulatory genes. 3T3 fibroblasts were treated with either non-targeting (NT) siRNA and DMSO (DMSO), NT siRNA and Cmpd 2 (CMPD2), or JNK1/2-targeting siRNA and Cmpd 2 (CMPD2+JNKsiRNA). All experimental groups were treated with TGF-β1 (2 ng/mL) for 24 hr. (A) Gene set enrichment analysis (GSEA) of TGF-β-target gene expression (KARLSSON_TGFB1_TARGETS_UP) comparing the CMPD2 treatment group to DMSO plus CMPD2+JNKsiRNA groups (REST). (Enrichment score = -0.6300272, Normalized Enrichment Score = - 2.7833443.) (B) Heatmap comparing expression of genes canonically upregulated or downregulated by TGF-β signaling between groups. (C) To identify MKP-5-JNK-regulated genes that mediate changes in TGF-β signaling, we identified targets that were either upregulated or downregulated by Cmpd 2-treatment (CMPD2 vs DMSO; 0.80 ≥ fold change ≥ 1.2, *p* ≤ 0.05) and rescued by JNK1/2 knockdown (CMPD2+JNKsiRNA vs CMPD2; 0.80 ≥ fold change ≥ 1.2, *p* ≤ 0.05). Candidate genes are represented within the overlapping area of the illustrated Venn diagrams. (D) Genes shown to activate TGF-β signaling, *Acta2, Thbs1, Inhba, Nuak2* and *Cdh11.* 3T3 fibroblasts were treated with NT/JNK1/2 siRNA, and DMSO/Cmpd 2. All groups were treated with TGF-β1 (2 ng/mL) for 24 hr and expression of genes was determined by qPCR.

## Discussion

In this study, we provide mechanistic insight into the role of MKP-5-mediated JNK regulation in TGF-β signaling thereby providing a plausible pathway through which MKP-5 promotes fibrosis. ^16,18,46^ We show that MKP-5 regulates TGF-β signaling by promoting TGF-β driven SMAD2 phosphorylation at canonical and non-canonical sites, nuclear translocation, and fibrogenic gene expression. We propose that MKP-5 mediates a JNK-dependent pathway that activates the expression of multiple TGF-β signaling activators (Figure 7). We found that in fibroblasts, TGF-β stimulation acutely upregulated MKP-5 expression (2 hr) and subsequently suppressed its expression at later timepoints (24 hr). This observation suggests a potential TGF-β feedback mechanism whereby TGF-β stimulation initially upregulates MKP-5 expression to promote signaling, but is later downregulated, to suppress TGF-β signaling. However, additional regulatory mechanisms are highly likely to be engaged by the MKP-5-JNK axis given the acute inhibitory effect that loss of MKP-5 function has on TGF-β-induced SMAD2 phosphorylation (Figure 7). This supposition is supported by two lines of orthogonal experimentation; MKP-5-deficient fibroblasts and pharmacological inactivation of MKP-5 activity both result in the failure of TGF-β to stimulate SMAD2 phosphorylation.

**Figure 7.**
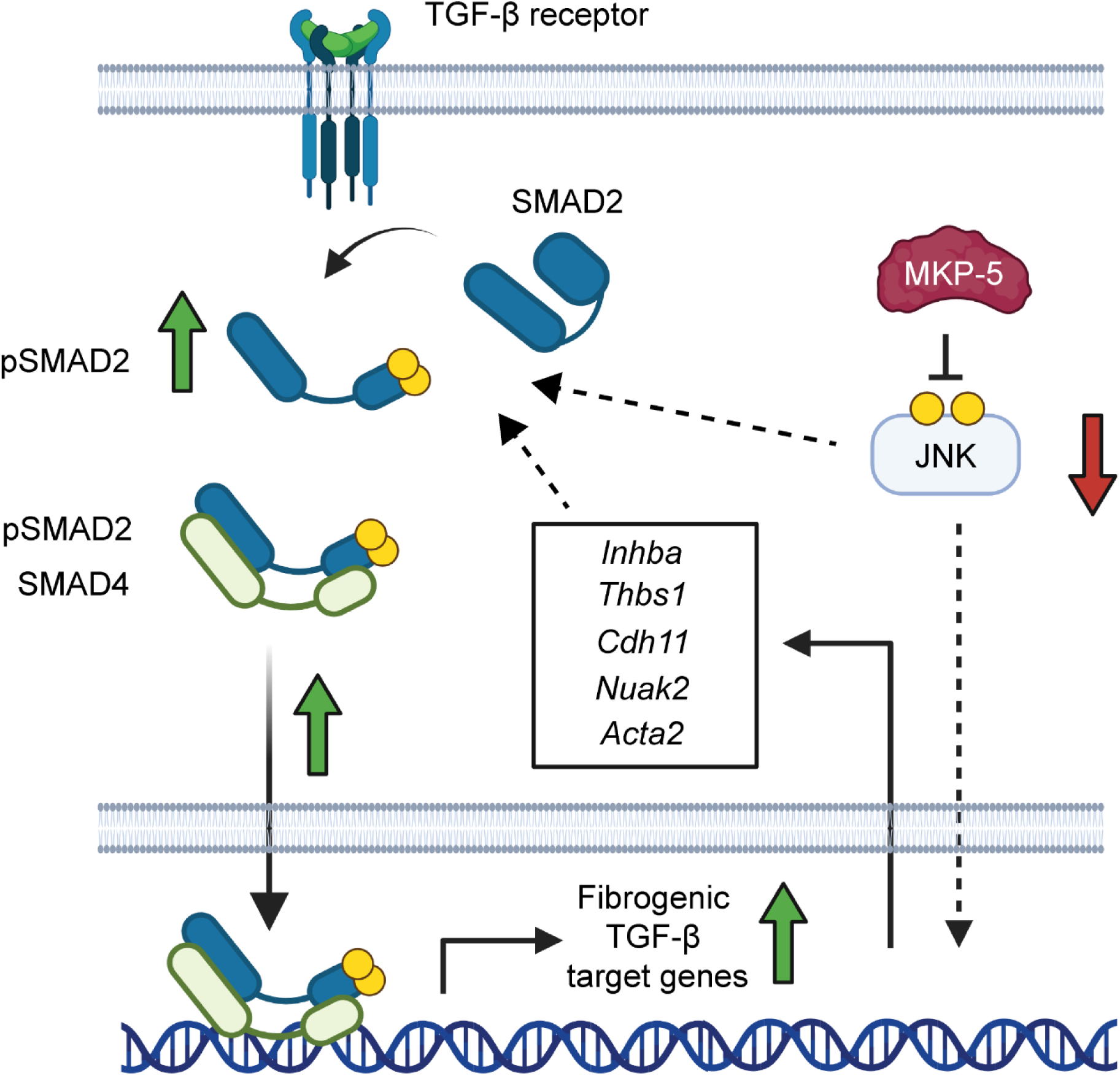
Model of MKP-5 regulating TGF-β signaling. In response to TGF-β, MKP-5 is required for both canonical and non-canonical TGF-β signaling in a JNK-dependent manner. In the canonical pathway, MKP-5-JNK activity regulates SMAD2 phosphorylation and nuclear translocation. In addition, MKP-5 regulates non-canonical TGF-β signaling. Following TGF-β activation MKP-5 inhibits JNK activity, which stimulates the expression of *Acta2, Thbs1, Inhba, Nuak2* and *Cdh11* genes that promote TGF-β activation.

We show that MKP-5 promotes TGF-β-driven phosphorylation of SMAD2 at multiple phosphorylation sites, including the c-terminus (S465/467) and the linker region (S245/250/255). C-terminus phosphorylation is mediated directly by the activated TGF-β receptor and is required for SMAD2 complex formation with SMAD4, followed by nuclear translocation.^47,48^ In contrast, the SMAD2 linker region is phosphorylated by other kinases, including p38 MAPK, JNK, ERK, glycogen synthase kinase, phosphatidylinositol 3’-kinase, Rho kinase, and cyclin-dependent kinases.^49–52^ The TGF-β receptor can indirectly activate many of these pathways, thus linker phosphorylation is often induced in response to TGF-β stimulation.^50,52,53^ However, the linker region of SMAD2 can also be phosphorylated in a TGF-β-independent manner.^38,54^ Although linker phosphorylation was initially characterized as an inhibitory mechanism, it has since been demonstrated that it can have either activating or inhibiting effects depending on the cellular context.^54–56^ Our results show that MKP-5 promotes both TGF-β-dependent c-terminus and linker phosphorylation providing direct evidence that MKP-5 is required for TGF-β receptor activation. These observations are consistent with the explanation for why both canonical and non-canonical pathways are inhibited in MKP-5-deficient cells. This effect could occur through a variety of mechanisms, including regulating receptor endocytosis, dimerization, phosphorylation or combinations thereof.^57^ These possibilities will require further investigation. However, the efficacy of the TGF-β receptor antibodies continues to be technically limiting. Additionally, the identification of the MKP-5-regulated kinase responsible for SMAD2 linker phosphorylation is currently unknown. Nonetheless, it is evident that neither p38 MAPK nor JNK are likely candidates for SMAD2 linker phosphorylation given that the activities of these MAPKs are elevated in the absence of MKP-5 expression.

Our results demonstrate that the catalytic activity of MKP-5 is required for TGF-β signaling. Fibroblasts treated with the MKP-5 inhibitor (Cmpd 2) suppressed TGF-β-dependent SMAD2 c-terminus and linker phosphorylation. Treatment of fibroblasts with Cmpd 2 resulted in the hyperactivation of MKP-5 substrates p38 MAPK and JNK, but not ERK, consistent with what is observed in MKP-5-deficient mice and cells.^17,35^ Given that SMAD2 c-terminus phosphorylation is required for its nuclear translocation and subsequent expression of fibrogenic genes, Cmpd 2 was also effective at inhibiting TGF-β-driven SMAD2 nuclear translocation and transcriptional activation of fibrogenic genes. Not surprisingly, the kinetics of SMAD2 phosphorylation (c-terminus and linker) differ between MKP-5-deficient fibroblasts and pharmacological inhibition of MKP-5 catalysis. TGF-β-stimulated MKP-5-deficient fibroblasts exhibited rapid inhibition of SMAD2 phosphorylation and nuclear translocation as compared with TGF-β-stimulated MKP5 inhibitor treated cells. It is likely that this difference is due to incomplete inhibition of MKP-5 phosphatase activity by Cmpd 2 and/or that MKP-5 has a non-catalytic contribution to regulating TGF-β signaling. Further studies will be required in order to parse out this differential putative role of catalytic and non-catalytic-mediated MKP-5 actions. It is noteworthy, that MKP-5 resides both within the nucleus and cytosol and it is conceivable that its sub-cellular localization might play a role in the catalytic and non-catalytic mechanisms of TGF-β action if operable.^58^

Direct inhibition of TGF-β signaling has been a long-standing common strategy for treating fibrosis.^59,60^ However, TGF-β has important physiological roles in immunoregulation, and therapeutic agents targeting often elicits severe side effects.^2,28,29^ These side effects have been ascribed to direct inhibition of TGF-β receptor kinase activity. Here, through inhibition of TGF-β signaling via MKP-5 antagonism the effects are likely to be TGF-β kinase-independent thereby raising the possibility that this mode of inhibition could yield less severe side effects. As such, the possibility of targeting MKP-5 for the treatment of fibrotic tissue disease is worthy of further experimental investigation. This is further bolstered by the observation that MKP-5-deficiency in various mouse models that develop tissue fibrosis in skeletal muscle, lung and the heart are all impaired. ^16,18,46^ It will be imperative to test MKP-5 inhibitors in these mouse models of tissue fibrosis. However, although Cmpd 2 is much improved over Cmpd 1 (Supplemental Table 1) it is not sufficiently efficacious to perform such *in vivo* studies and additional development of these MKP-5 inhibitors is required for these studies to be performed.

Mechanistically, our results have uncovered a novel regulatory mechanism of TGF-β signaling. Although the precise mechanism of how MKP-5 is directly involved in TGF-β-mediated SMAD2 phosphorylation currently remain elusive, regardless our data implicate the actions of an MKP-5-JNK-dependent axis. MKP-5 has been shown to regulate cellular function by limiting the activity of its MAPK substrates, p38 MAPK and JNK, via dephosphorylation. For this reason, we suspected that when hyperactivated as a result of MKP-5 inhibition, subsequent restoration of either p38 MAPK and/or JNK would rescue TGF-β signaling in Cmpd 2 treated cells. Our results showed that inhibition of neither p38 MAPK nor ERK rescued TGF-β signaling. The lack of effect with ERK was not surprising given that ERK is a very poor MKP-5 substrate, but p38 MAPK appeared to accentuate the inhibition suggesting another role for MKP-5-p38 MAPK in this pathway that remains to be resolved. Importantly, both pharmacologic inhibition and siRNA knockdown of JNK rescued TGF-β signaling in Cmpd 2-treated cells. This demonstrates that MKP-5 regulates TGF-β through a JNK-dependent pathway. Previous reports investigating the role of JNK in TGF-β signaling have produced varying conclusions. In one study investigating the role of JNK signaling in lung development, mouse embryonic fibroblasts treated with SP600125 were shown to have enhanced TGF-β-mediated SMAD2 c-terminus phosphorylation, as well as downstream transcription, consistent with our findings.^61^ However, studies in non-fibroblast cell lines have shown differential regulation. A study investigating TGF-β signaling in HepG2 cells found that SP600125 blocked TGF-β driven SMAD2 phosphorylation, both on the c-terminus and linker phosphorylation sites.^62^ Another group showed that myoblasts transfected with a constitutively active MKK7-JNK fusion protein blocked SMAD2 phosphorylation and nuclear translocation in response to myostatin stimulation.^38^ These data suggest that the role of JNK as a regulator of TGF-β signaling may vary depending on the cell type and at least in fibroblasts it appears to negatively regulate TGF-β signaling.

We tested the hypothesis that the MKP-5-JNK signaling axis may regulate the TGF-β pathway by modulating expression of TGF-β regulatory genes. To identify which genes may be involved, we performed RNA sequencing on Cmpd 2-treated fibroblasts with or without JNK1/2 knockdown to isolate relevant targets. From our analysis, we identified a subset of TGF-β activators that were downregulated by Cmpd 2 and rescued by JNK knockdown, consistent with the observed effect on TGF-β signaling. Notable from this list is *Thbs1*, a gene that is involved with TGF-β ligand processing and promotes an autocrine, positive feedback loop.^43^ *Cdh11* promotes TGF-β expression, also promoting autocrine signaling.^44^ *Inhba* is a subunit of the Activin ligand, which promotes SMAD2 phosphorylation and synergizes with TGF-β signaling.^63^ *Nuak2* binds R-SMADs and prevents their degradation.^45^ It is conceivable that at least one mechanism through which MKP-5-JNK regulates TGF-β signaling is through the transcriptional regulation of these signaling activators. It is also possible that MKP-5-JNK regulates TGF-β signaling through non-transcriptional mechanisms as alluded to earlier. For example, in the absence of MKP-5, JNK hyperactivation could drive phosphorylation of substrates that regulate other processes such as TGF-β receptor endocytosis, dimerization, or R-SMAD localization. Future phosphoproteomic analysis on MKP-5 deficient and/or Cmpd 2-treated cells will be necessary to identify substrates regulated by JNK directly, which would elucidate these additional potential mechanisms.

Multiple other protein tyrosine phosphatases have been shown to be involved in TGF-β signaling. For example, DUSP8 regulates expression of *Il9* in response to TGF-β signaling by dephosphorylating the transcriptional repressor Pur-α.^64^ SHP2 has been shown to regulate non-canonical TGF-β-driven STAT3 activation by dephosphorylating JAK2 in response to stimulation.^65^ PTP1-α has been shown to promote pulmonary fibrosis and TGF-β driven SMAD2 phosphorylation through a Src-dependent mechanism.^66^ Notably, MKP-2 was shown to inhibit TGF-β-driven SMAD2 phosphorylation through a JNK-dependent mechanism, seemingly demonstrating the opposite function of MKP-5.^67^ This difference is likely due to the role of MKP-2 being measured within NRK-52E cells as opposed to fibroblasts. Nevertheless, these examples illustrate the complexities of TGF-β regulation. Nonetheless, the data presented here coupled with our previous studies on the actions of MKP-5 in tissue fibrosis provide strong evidence that MKP-5 is intimately involved in maintaining tissue homeostasis through TGF-β regulation.

## Methods

### Cell culture and luciferase reporter cell assay

Cell-based assays used 3T3 fibroblasts, mouse embryonic fibroblasts (MEFs), or A549 cells grown at 37 °C and 5% CO_2_. *Mkp5^+/+^* and *Mkp5^−/−^* mouse embryo fibroblasts were generated from female mice at days 13-14 of pregnancy and established by spontaneous immortalization. *Mkp5^-/-^*mice were provided by Dr. Flavell (Yale University School of Medicine) and were maintained as previously described.^17,68^ Cells were maintained in Dulbecco’s modified eagle’s medium (DMEM) (11965-092, Gibco) supplemented with 10% fetal bovine serum (FBS) (F0926, Sigma Aldrich), 1% sodium pyruvate (11360-70, Gibco), and 1% penicillin-streptomycin (15140-122, Gibco).

The TGF-β reporter construct was generated by inserting four Smad-binding-element repeats (GTCTAGAC) into a PGL4.2 expression vector. A549 cells were stably transfected with the reporter construct and transfected clones were selected with 1 ug/mL puromycin. 5 x 10^5^ cells were plated on 12-well plates and treated with Cmpd 2 (10 µM), MAPK inhibitors (10 µM), and/or TGF-β1 (2 ng/mL) for 24 hr. MAPK inhibitors U0126 (9903), SP600125 (8177), and SB203580 (5633) were purchased from Cell Signaling Technologies. TGF-β1 (240-B-002) was purchased from R&D Systems. Cells were lysed, mixed with luciferase reagent, incubated for two minutes, and luminosity was measured as previously described.^69^

### Quantitative PCR analysis

RNA was isolated from cells using QIAGEN RNeasy kit. 1 µg RNA was reverse transcribed to generate cDNA using a reverse transcriptase PCR kit. Real-time quantitative PCR was performed using 7500 Fast real-time PCR system, and either Power UP SYBR Green Master Mix or Taqman Master Mix kit. Relative gene expression was calculated using ΔΔCt method and was normalized to either 18S rRNA or GAPDH expression. Primers used for Taqman qPCR: *Col1a1* (Mm00801666), *Fn1* (Mm00725412), *Acta2* (Mm00725412), *Gapdh* (Mm99999915), and *18s* (Hs99999901) were purchased from Thermo Fisher Scientific. Primers used for SYBR Green qPCR: *18s* (F: GGCCTCACTAAACCATCCAA, R: GCAATTATTCCCCATGAACG), *Dusp10* (F: CCATCTCCTTTAGACGACAGGG, R: GCTACCACTACCTGGGCTG), *Nuak2* (F: ATCAAGTCGCCTAAACCTCTGA, R: CCTCCGTATGTGCAGCAGAT), *Cdh11* (F: CTGGGTCTGGAACCAATTCTTT, R: GCCTGAGCCATCAGTGTGTA), *Thbs1* (F: GGGGAGATAACGGTGTGTTTG, R:CGGGGATCAGGTTGGCATT), *Inhba* (F: TCAGAGGATTTCTGTTGGCAAG, R: TCACATCGGGTCTCTTCTTCA).

### Immunoblotting

Cells were lysed on ice in lysis buffer (50mM Tris HCl, 150mM NaCl, 5mM EDTA, 1% NP-40, 0.5% SDS, 0.5% deoxycholate acid). Cells were incubated on ice for 30 minutes, then debris was pelleted using centrifugation. Protein concentration was measured by BCA analysis (23225, Thermo Fisher Scientific). Cells lysates were resolved by SDS-PAGE, transferred to nitrocellulose membrane, blocked with 5% BSA (A9647, Sigma-Aldrich), and immunoblotted. Antibody binding was visualized using the Odyssey CLx Imaging System (LI-COR Biosciences). The following primary antibodies were used: total SMAD2 (3103), phospho-SMAD2 (S465/467) (3108), phospho-SMAD2 (S245/250/255) (3104), phospho-p38 MAPK (T180/Y182) (9215), phospho-JNK (T183/185) (4668), total JNK (3708), phospho-ERK (T202/Y204) (3101), and total ERK (9107), purchased from Cell Signaling Technologies; total p38α MAPK (81621) and GAPDH (13719) antibodies were purchased from Santa Cruz Biotechnology; α smooth muscle Actin (5694) antibody was purchased from Abcam.

### Immunofluorescence and confocal microscopy

*Mkp5^+/+^* and *Mkp5^−/−^* mouse embryo fibroblasts or 3T3 fibroblasts (5 x10^5^) were seeded in growth media onto coverslips. MEFs were serum starved (0.1% FBS in DMEM) for 4 hr, then stimulated with TGF-β1 (2 ng/mL) for 10 minutes. 3T3 fibroblasts were treated with 10 µM Cmpd2 or DMSO in 0.1% FBS in DMEM for 4 hr, then stimulated with TGF-β (2 ng/mL) for 1 hr. Cells were fixed in 4% paraformaldehyde, permeabilized with 0.1% Triton X-100, then incubated with SMAD2 antibody (5339, Cell Signal Technologies) at 4 °C overnight. Cells were washed and incubated with secondary antibody (AlexaFluor 488 Goat-anti-rabbit; A11008 Thermo Fisher Scientific) and anti-phalloidin conjugated antibody (AlexaFluor 594; A12381 Thermo Fisher Scientific) for 2 hr at room temperature. Cells were mounted in DAPI mounting media (ab104139, Abcam). Microscopy was performed on a Zeiss Airyscan using ZEN Black software. SMAD2 intensity was quantified using ImageJ.

### RNA sequencing analyses

RNA was isolated cells using the QIAGEN RNeasy kit and RNA quality was determined by estimating the A260/A280 and A260/A230 ratios by NanoDrop (Thermo Fisher Scientific). RNA integrity was determined by running an Agilent Bioanalyzer gel, which measures the ratio of the ribosomal peaks. mRNA was purified from approximately 50 ng of total RNA with oligo-dT beads and sheared by incubation at 94 °C in the presence of Mg^2+^ (KAPA mRNA HyperPrep). Following first-strand synthesis with random primers, second strand synthesis and A-tailing were performed with dUTP to generate strand-specific sequencing libraries. Adapter ligation with 3′-dTMP overhangs were ligated to library insert fragments. Library fragments carrying the appropriate adapter sequences at both ends were amplified using high-fidelity, low-bias PCR. Strands marked with dUTP were not amplified. Indexed libraries that met appropriate cutoffs for both were quantified by qRT-PCR using a commercially available kit (KAPA Biosystems) and insert size distribution was determined with the LabChip GX or Agilent Bioanalyzer. Samples with a yield of at least 0.5 ng/μL were used for sequencing. Sample concentrations were normalized to 1.2 nM and loaded onto an Illumina NovaSeq flow cell at a concentration that yielded 25 million passing filter clusters per sample. Samples were sequenced using 100-bp paired-end sequencing on an Illumina NovaSeq according to Illumina protocols. The 10-bp dual index was read during additional sequencing reads that automatically followed the completion of read 1. Data generated during sequencing runs were simultaneously transferred to the Yale Center for Genome Analysis (YCGA) high-performance computing cluster. A positive control (prepared bacteriophage Phi X library) provided by Illumina was spiked into every lane at a concentration of 0.3% to monitor sequencing quality in real time. Signal intensities were converted to individual base calls during a run using the system’s Real Time Analysis software. Base calls were transferred from the machine’s dedicated personal computer to the Yale high-performance computing cluster via a 1 Gigabit network mount for downstream analysis.

### Pathway Analysis

The RNA-Seq statistical analysis was performed using Partek Flow Genomic Analysis software build version 8.0.19.1125 (Partek Inc.). Paired-end reads were trimmed and aligned to the Genome Reference Consortium Mouse Build 38 (mm10) with the STAR alignment tool (ver. 2.6.1d). Total counts per gene were quantified and normalized to identify differentially expressed genes (DEGs). Lists of DEGs or transcripts were generated by DESeq2. Qlucore Omics Explorer 3.5 (Qlucore AB) was used for principal component analysis of log_2_-transformed global expression values, heatmap generation, and hierarchical clustering. DEGs used in pathway analysis were determined by using a filtering criterion of fold change ≤ –1.2 or ≥ 1.2 (*p* ≤ 0.05), Benjamini-Hochberg multiple testing correction *p*-value. Ingenuity Pathway Analysis (QIAGEN) software (ver. 10-14) was used to identify top upstream regulators, top canonical pathways, diseases, and functions overrepresented in the DEGs.

### Cell Viability Assays

Cell viability was measured using the MTS Assay (Abcam, ab197010) and Trypan Blue (ThermoFisher Scientific, 15250061). 3T3 fibroblasts (5 x 10^5^) were cultured in DMEM growth media and cultured for 24 hr in various concentrations of Cmpd 2, ranging from 0-30 µM. MTS Assay was performed according to the manufacturer’s listed protocol. For Trypan Blue Assay, cells were combined with 0.4% Trypan Blue solution in a 1:1 ratio, and viable cells were counted using a hematocytometer.

### Statistical analyses

No statistical methods were used to predetermine sample size. The number of samples used in each experiment is shown. All *in vitro* experiments were performed at least three times independently. Statistical analysis and graphing were performed using GraphPad Prism 10.0.2 software. All data represent the means ± SEM. For *p* value determinations, we used one-way or two-way ANOVA with multiple comparisons.

## Supporting information

Dorry et al_Supplemental figures and tables

## Author Contributions

S.J.D. and A.M.B. were responsible for the conceptual design of all the experiments. S.J.D. performed all the experiments. S.P. assisted in the analysis of whole genome RNA sequencing data. All data were analyzed and interpreted by S.J.D. and A.M.B. The manuscript was written by S.J.D. and A.M.B.

## Disclosure statement

A.M.B. is founder of Allagium Therapeutics. All the other authors declare no competing financial interests.

## Data availability

All data needed to evaluate the conclusions in the paper are present in the paper and/or the Supplementary Materials.

## Funding

This works was supported by NIH grants R01 AR080152 and R01 HL158876 to A.M.B. S.J.D. received support from the pre-doctoral Pharmacology Training Program (T32 GM0073241). We thank Lei Zhang for technical assistance.

