## Supplementary material for "MAPK Phosphatase-5 is required for TGF-β signaling through a JNK-dependent pathway": Dorry et al_Supplemental figures and tables

Supplemental Figure 1. Effect of Cmpd 2 on cell proliferation or viability

Supplemental Figure 2. Effect of Cmpd 2 on TGF-β-mediated gene expression in *Mkp5*$^{-/-}$ fibroblasts

Supplemental Figure 3. p38 MAPK inhibition does not rescue TGF-β-target gene expression in Cmpd 2-treated fibroblasts

Supplemental Figure 4. Principal Component Analysis of RNA sequencing samples

Supplemental Table 1. Characterization of Cmpd 2 as compared with Cmpd 1

Supplemental Table 2. Cmpd 2 inhibits expression of fibrosis-related genes

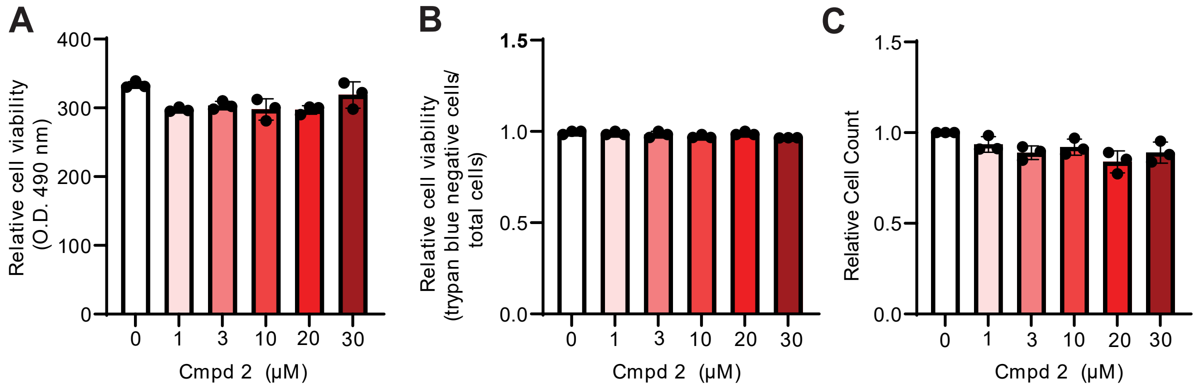


**Figure S1.** Effect of Cmpd 2 on cell proliferation and cell viability. 3T3 fibroblasts were treated with the indicated concentrations of Cmpd 2 for 24 hr, then assessed with (A) MTS Assay or (B) Trypan blue assay to determine cell viability. (C) Cells were treated with the indicated concentration of Cmpd 2 for 24 hr, then cells were counted to determine the effect of Cmpd 2 on proliferation. Data represent the mean ± SEM derived from 3 independent experiments.

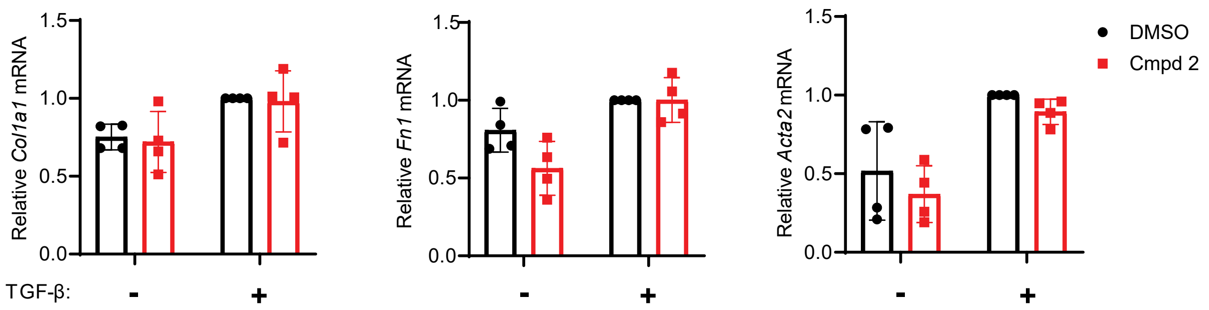


**Figure S2.** Effect of Cmpd 2 on TGF-β-mediated gene expression in *Mkp5*$^{-/-}$ fibroblasts. *Mkp5*$^{-/-}$ fibroblasts were treated with Cmpd 2 (10 μM) and TGF-β (2ng/mL) overnight. Relative expression of TGF-β-target genes *Col1a1, Fn1,* and *Acta2* was measured using qPCR. Data represent the mean ± SEM derived from 4 independent experiments.

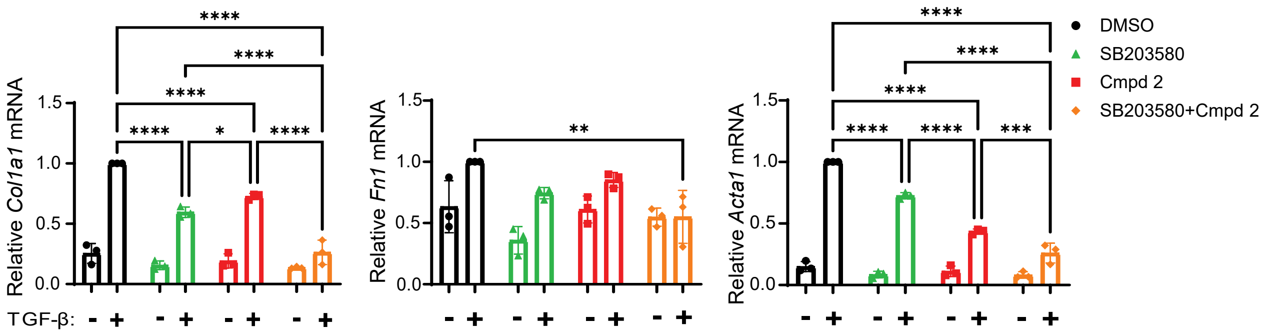


**Figure S3.** p38 MAPK inhibition does not rescue TGF-β-target gene expression in Cmpd 2-treated fibroblasts. NIH3T3 fibroblasts were treated with p38 MAPK inhibitor (SB203580, 10 µM), Cmpd 2 (10 µM), and/or TGF-β (2 ng/mL) for 24 hr. Expression of TGF-β-target genes *Col1a1, Fn1,* and *Acta2* was measured using qPCR. Data represent the mean ± SEM derived from 3 independent experiments.

Key: **: *p*-value<0.01, ***: *p*-value<0.001, ****: *p*-value<0.0001.

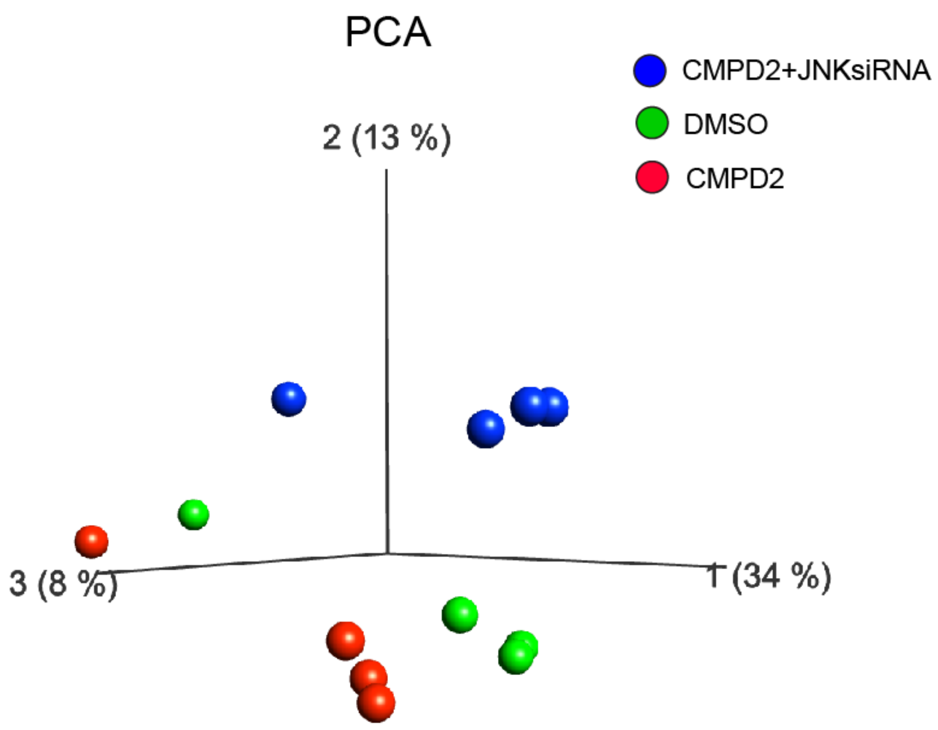


**Figure S4.** Principal Component Analysis of RNA sequencing samples. 3T3 fibroblasts were treated with either non-targeting (NT) siRNA and DMSO (DMSO), NT siRNA and Cmpd 2 (CMPD2), or JNK1/2-targeting siRNA and Cmpd 2 (CMPD2+JNKsiRNA). All experimental groups were treated with TGF-β1 (2 ng/mL) for 24 hr. Total RNA was isolated and RNA sequencing was performed. Principal Component Analysis of the three experimental groups is illustrated.

| | Compound 1 | Compound 2 |
|---|---|---|
| MKP-5 catalytic domain inhibition [$IC_{50}$ (μM)] | 3.9 | 0.2 |
| Selectivity [MKP-5/MKP-7/MKP-1; $IC_{50}$ (μM)] | 2.2/ >35/ >35 | 1.5/ 23/ 35 |
| Cellular pp38 MAPK/p38 MAPK [Fold increase, 10μM] | ~3 | ~7 |

**Table S1.** Characterization of Cmpd 2 as compared with Cmpd 1. Table comparing the potency and selectivity of MKP-5 inhibitors, Compound 1 (Cmpd 1) and Compound 2 (Cmpd 2).

| Gene name | Expr p-value | Expr Fold Change (CMPD2 vs DMSO) | ID | Predicted relationship |
|---|---|---|---|---|
| *Aspn* | 2.13E-02 | -1.292 | ENSMUSG00000021388 | not predicted |
| *Atp2b4* | 8.35E-02 | -1.337 | ENSMUSG00000026463 | not predicted |
| *Bdkrb1* | 1.21E-05 | 1.406 | ENSMUSG00000021070 | not predicted |
| *Bdkrb2* | 3.98E-03 | 1.368 | ENSMUSG00000021070 | inhibits |
| *Ccn2* | 3.22E-04 | -1.337 | ENSMUSG00000019997 | inhibits |
| *Cdc42ep3* | 3.28E-03 | -1.253 | ENSMUSG00000036533 | not predicted |
| *Cdh11* | 3.60E-03 | -1.467 | ENSMUSG00000031673 | inhibits |
| *Chi3l1* | 8.51E-15 | 1.986 | ENSMUSG00000064246 | activates |
| *Clu* | 2.89E-02 | -1.38 | ENSMUSG00000022037 | activates |
| *Cyp1b1* | 9.02E-04 | 1.618 | ENSMUSG00000024087 | activates |
| *Edn1* | 1.76E-02 | -1.522 | ENSMUSG00000021367 | not predicted |
| *Grin2d* | 7.01E-02 | -1.264 | ENSMUSG00000002771 | not predicted |
| *Igf1* | 6.83E-06 | -1.419 | ENSMUSG00000020053 | activates |
| *Inhba* | 3.36E-05 | -1.69 | ENSMUSG00000041324 | inhibits |
| *Irf5* | 4.18E-02 | -1.262 | ENSMUSG00000029771 | inhibits |
| *Jak3* | 9.43E-02 | 1.271 | ENSMUSG00000031805 | not predicted |
| *Krt8* | 6.84E-02 | -2.09 | ENSMUSG00000049382 | inhibits |
| *Lpl* | 2.74E-02 | -1.519 | ENSMUSG00000015568 | activates |
| *Maf* | 1.57E-03 | 1.452 | ENSMUSG00000055435 | not predicted |
| *Mfsd2a* | 6.85E-03 | -1.45 | ENSMUSG00000028655 | activates |
| *Mmp13* | 9.14E-02 | -1.555 | ENSMUSG00000050578 | inhibits |
| *Mmp9* | 3.16E-03 | -1.341 | ENSMUSG00000017737 | activates |
| *Ramp3* | 1.34E-02 | 1.26 | ENSMUSG00000041046 | inhibits |
| *Rgs5* | 1.57E-02 | -1.856 | ENSMUSG00000026678 | activates |
| *Scx* | 8.30E-02 | -1.647 | ENSMUSG00000034161 | not predicted |
| *Serpina3* | 4.77E-03 | 1.265 | ENSMUSG00000021091 | inhibits |
| *Ssc5d* | 2.22E-03 | -1.448 | ENSMUSG00000035279 | not predicted |
| *Thbs1* | 4.95E-06 | -1.523 | ENSMUSG00000040152 | activates |
| *Tnc* | 8.88E-03 | -1.292 | ENSMUSG00000028364 | inhibits |
| *Tnfrsf11b* | 7.44E-02 | -1.527 | ENSMUSG00000063727 | inhibits |
| *Tnnt2* | 1.79E-07 | -1.35 | ENSMUSG00000026414 | not predicted |
| *Usp18* | 1.84E-02 | -1.562 | ENSMUSG00000030107 | activates |
| *Xdh* | 1.29E-08 | 1.495 | ENSMUSG00000024066 | inhibits |
| *Acta2* | 1.64E-05 | -1.42 | ENSMUSG00000035783 | inhibits |
| *Angpt1* | 4.12E-03 | -3.107 | ENSMUSG00000022309 | not predicted |
| *Apob* | 2.44E-02 | -1.619 | ENSMUSG00000020609 | not predicted |

**Table S2.** Cmpd 2 inhibits expression of fibrosis-related genes. RNA sequencing was performed on NIH3T3 cells treated with TGF-β and Cmpd 2 or DMSO overnight. Ingenuity Pathway Analysis was performed to identify DEGs associated with various diseases, including fibrosis.
